# plasmoRUtils: A one-stop R Package for Plasmodium and other Apicomplexan parasite-related Bioinformatics analysis

**DOI:** 10.1101/2025.07.30.667718

**Authors:** Rohit Satyam, Alberto Maillo, Amit Kumar Subudhi, Rahul P Salunke, David Gomez-Cabrero, Arnab Pain

## Abstract

Bioinformatics analysis of non-model organisms remains challenging due to the limited availability of specialized tools, as most R packages are optimized for well-annotated model species. This problem is exacerbated by genomic and proteomic data being scattered across multiple databases, each employing different identifiers based on varying reference annotations. Comprehensive databases have been developed to disseminate knowledge related to Apicomplexan genomics and proteomics, such as VEupathDB. Several specialised databases, particularly for the malaria parasite *Plasmodium*, have been developed such as ApicoTFDB, malaria.tools, MPMP, MIIP, Phenoplasm, PlasmoBase, alongside broader resources like HitPredict and TED. However, these platforms often suffer from manual query interfaces, outdated identifiers, and inefficient data retrieval methods, complicating their use for large-scale bioinformatic analyses. To address these limitations, we present plasmoRUtils, an R package designed to streamline database access and data harmonization for Apicomplexan research. plasmoRUtils enables the retrieval of data tables using single-line R functions and standardized Ensembl gene IDs as input. Thanks to the APIs available for some databases such as VEuPathDB, the package also provides functions to build CLI-based queries for VEuPathDB’s component databases. Additional support includes performing overrepresentation analyses and estimating parasite transcriptomic age/stage using single-cell or bulk RNA-Seq references. By automating data retrieval and transformation within RStudio, plasmoRUtils eliminates the need for manual database queries, facilitating end-to-end workflow development without leaving the R environment. Available on GitHub https://github.com/Rohit-Satyam/plasmoRUtils, with comprehensive documentation, plasmoRUtils represents a critical step toward efficient and reproducible bioinformatics for Apicomplexan research.

## 1. Introduction

Within the R ecosystem, conducting bioinformatic analyses of non-model organisms poses significant challenges. Most widely used R packages and computational pipelines have been developed and validated primarily using data from model organisms, limiting their applicability to less well-annotated species. This issue is further complicated by the fact that most of the data for non-model organisms is often scattered across multiple specialized and privately maintained databases or shiny apps, each using distinct gene or protein identifiers, based on varying genome assemblies and annotations available at the time of their development. Besides, these databases are not synchronously updated. This inconsistency complicates tasks such as gene mapping, cross-referencing orthologs, and integrating multiple datasets without reanalysis (1).

To mitigate some of these issues in the context of Apicomplexan parasites—a diverse group of human and animal pathogens that includes *Plasmodium* spp., *Toxoplasma gondii*, *Cryptosporidium* spp., etc— a compendium of eukaryotic pathogen related resources called VEuPathDB (Eukaryotic Pathogen, Vector, and Host Informatics Resource) has been developed. The is a comprehensive bioinformatics resource that interfaces with 12 specialized databases for host and pathogen information. For two decades, it has provided user-friendly tools, dataset access, and up-to-date annotations for the global scientific community (2,3). It offers annotations and datasets for over 600 organisms, including invertebrate vectors, eukaryotic pathogens (such as protists and fungi), and relevant free-living or non-pathogenic species and hosts. VEuPathDB has significantly contributed to advancing research on various tropical diseases (4,5). In parallel with VEuPathDB, several privately maintained databases have emerged in recent years. In the field of malaria, these include malaria.tools (6), MPMP (7), ApicoTFDB (8), MIIP (9), and Phenoplasm (10), which host valuable predicted and curated information. Other non-pathogen-focused databases that host apicomplexan data include HitPredict (11,12), which provides *Plasmodium* protein-protein interaction data, and The Encyclopedia of Domains (TED) (13), which offers protein domain boundary information for both *Toxoplasma* and *Plasmodium*.

While some of the aforementioned biological databases offer facility for exporting underlying data in bulk, some static databases allow only single-query operations at a time. For example, VEuPathDB and its 12-component organism-specific databases (henceforth referred to as "VEuPathDBs") provide an online platform for conducting bioinformatics tasks such as ID mapping, enrichment analysis, and manual downloads of preconfigured tables (2). Additionally, they offer programmatic access through an Application Programming Interface (API) using the Representational State Transfer (REST) protocol, which is simpler, faster, and widely used for batch retrieval. On the contrary, the MPMP database contains the most detailed, manually curated gene sets for key malaria metabolic pathways derived from the literature (7) but require manual query and batch query or API is not available. Several published studies have conducted MPMP enrichment analyses in the past (14–18) but did not provide a snapshot of the database or the gene sets used at the time of their analysis. Since pathway maps and tables in MPMP are subject to periodic changes through manual curation, it is not possible to track which changes were made or which genes were added or removed from the gene sets. Some authors rely on MPMP data from PlasmoDB (19,20), which is stable at version 2019 (personally requested from Prof. Hagai Ginsburg) but has not been updated since. Further, a manual effort is often required in databases where users must navigate through multiple URLs to access the relevant tables. A case in point is the HitPredict database (11), which provides protein-protein interactions (PPIs) for 125 species, including *Plasmodium*, but only allows querying one UniProt ID at a time. Additionally, these protein IDs must be converted back to gene IDs before they can be used in genomic investigations, such as pathway enrichment analysis using *pathfindR* (21) or while combining co-expression analysis with PPI information. Other databases, such as MIIP (9) and Phenoplasm (10) (which contains outdated mitochondrial gene IDs and missing *Plasmodium* genes), use old gene IDs that complicate the quick and direct application of the data for hypothesis generation and analysis.

Previous efforts have been made to provide programmatic access to VEuPathDBs (formerly known as EuPathDB, https://github.com/khughitt/EuPathDB) through an R package. However, the EuPathDB R package is no longer maintained, and its functionality has largely deteriorated, rendering the package ineffective. Currently, no other R package offers practical access to VEuPathDBs data. For specific bioinformatics tasks, such as converting Gene IDs to UniProt IDs and vice versa, we attempted to use the *biomaRt* R package, which works well for well-studied pathogens like *Toxoplasma gondii* and *Plasmodium falciparum* (22) but has been observed not to work for all other pathogens such as *Plasmodium berghei* (Pb) (https://github.com/grimbough/biomaRt/issues/110). For Pb, the Protists Ensembl release 56, utilized the same reference genome assembly as used by PlasmoDB, however, the gene annotations were not updated since 2015 (PBANKA01, INSDC Assembly GCA_900002375.1, November 2015). In order to carry out ID conversion programmatically, the users must wait for Protists Ensembl to switch their annotation from ENA to VEuPathDB and synchronize it so that they can be used through biomaRt R package. The aforementioned issues prompted us to design a solution in the form of an R package, *plasmoRUtils*, which provides a more idiomatic and user-friendly method for programmatically accessing VEuPathDBs data.

Using *plasmoRUtils*, users can fetch data from VEuPathDBs and other databases and seamlessly transform it into formats compatible with downstream R packages. Leveraging the RESTful API provided by VEuPathDB, the package includes intuitive R functions for query construction, enabling the retrieval of both preconfigured and user-defined data tables directly within the R environment. For databases that do not offer programmatic access via APIs, we developed database-specific "searchX" functions (where X represents the database) that utilize the *rvest* package for web crawling to retrieve data, which is then transformed into tables that can be saved and shared (23,24). Additionally, we created a function to enable programmatic access to the MPMP database for the first time, allowing users to download and share data tables efficiently. The package also provides several other datasets that we reanalyzed using the latest annotations from VEuPathDBs, which various functions can utilize. We chose R as the development platform due to its widespread adoption in the bioinformatics community, its robust statistical capabilities, and the rich ecosystem of packages that support data transformation, visualization, and analysis—all of which can be readily applied to outputs generated by plasmoRUtils.

## 2. Materials and methods

For VEuPathDBs, we provide simplified access to the API within the R environment by developing wrapper "get" functions (see functions in Table 1 and Figure 1). This abstraction of the server interface eliminates the need to understand the database structure or the specific URL endpoints required to retrieve the necessary data. To facilitate access to non-API databases, we also offer various "search" functions. Some of these functions are specific to *Plasmodium* or related species; if the database only provides data and annotations related to *Plasmodium*, while others are species-agnostic. A detailed list of these functions, along with the relevant organisms and the rationale behind their development, is provided below and documented in the package vignette. Given a set of gene IDs to query, these functions perform web scraping of the databases and return the transformed data in the form of data tables, which can then be easily saved and shared.

**Figure 1:**
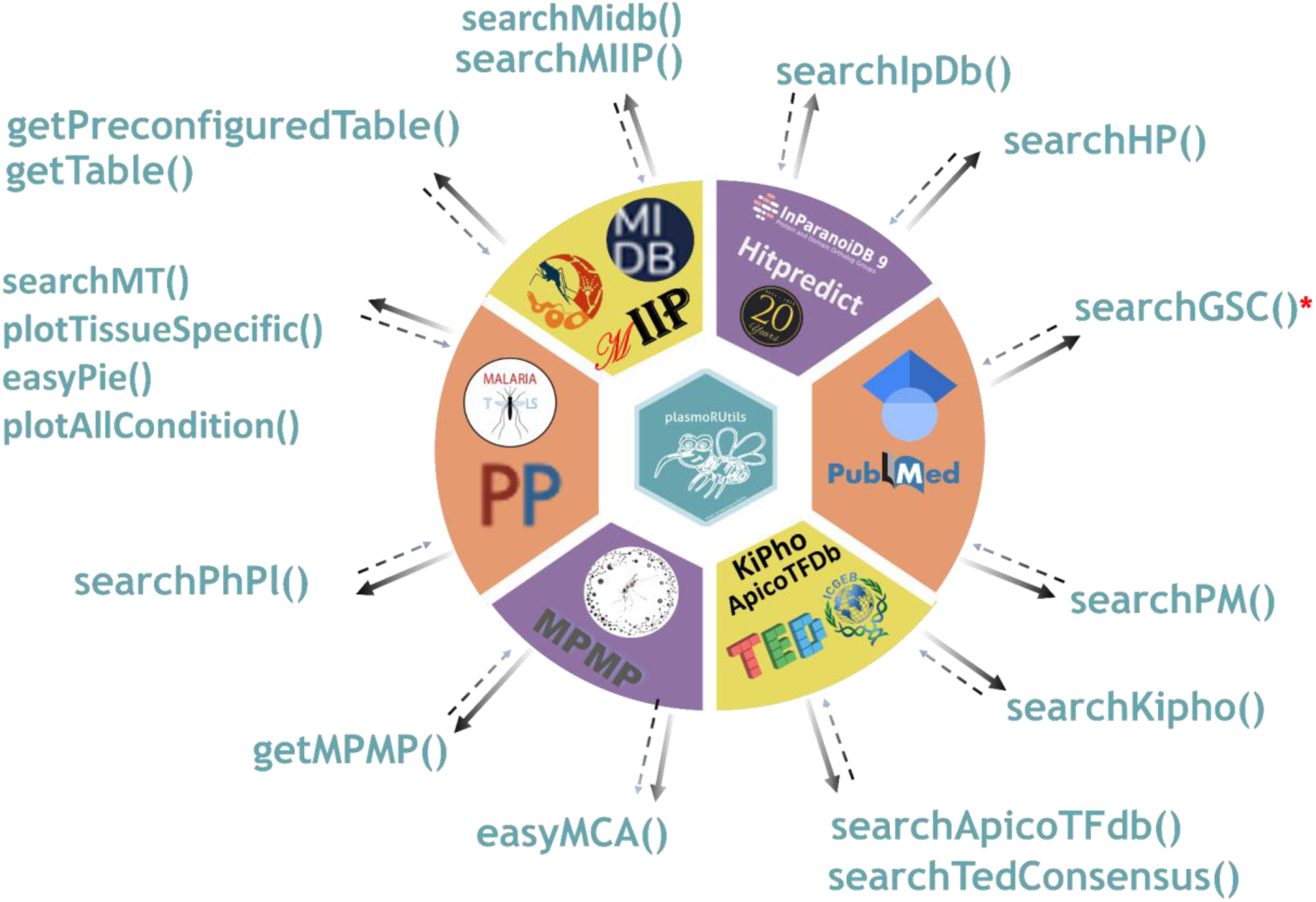
A graphical overview of the functions and database access available through the *plasmoRUtils* R package. *Limited queries per day.

**Table 1:**
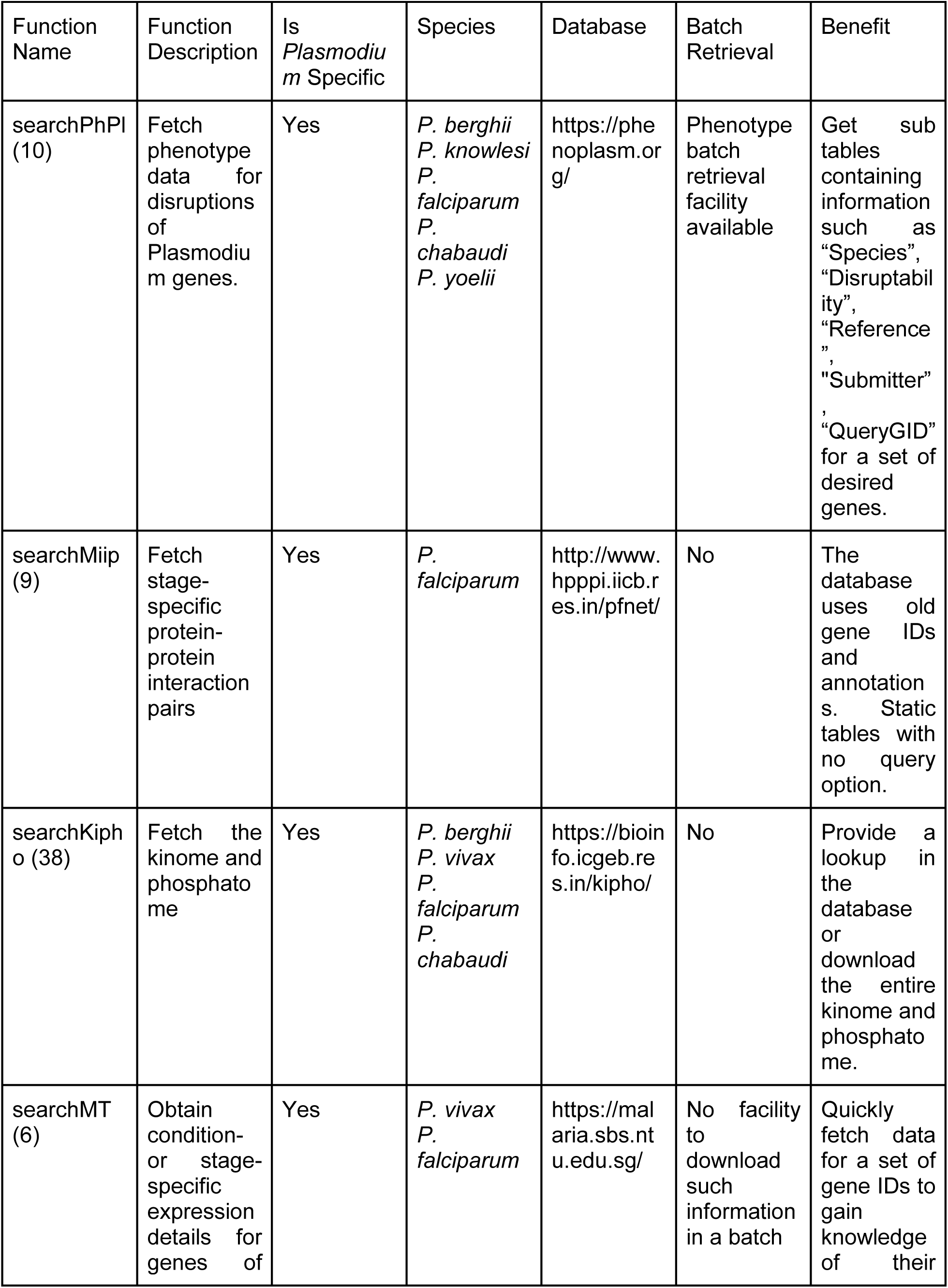
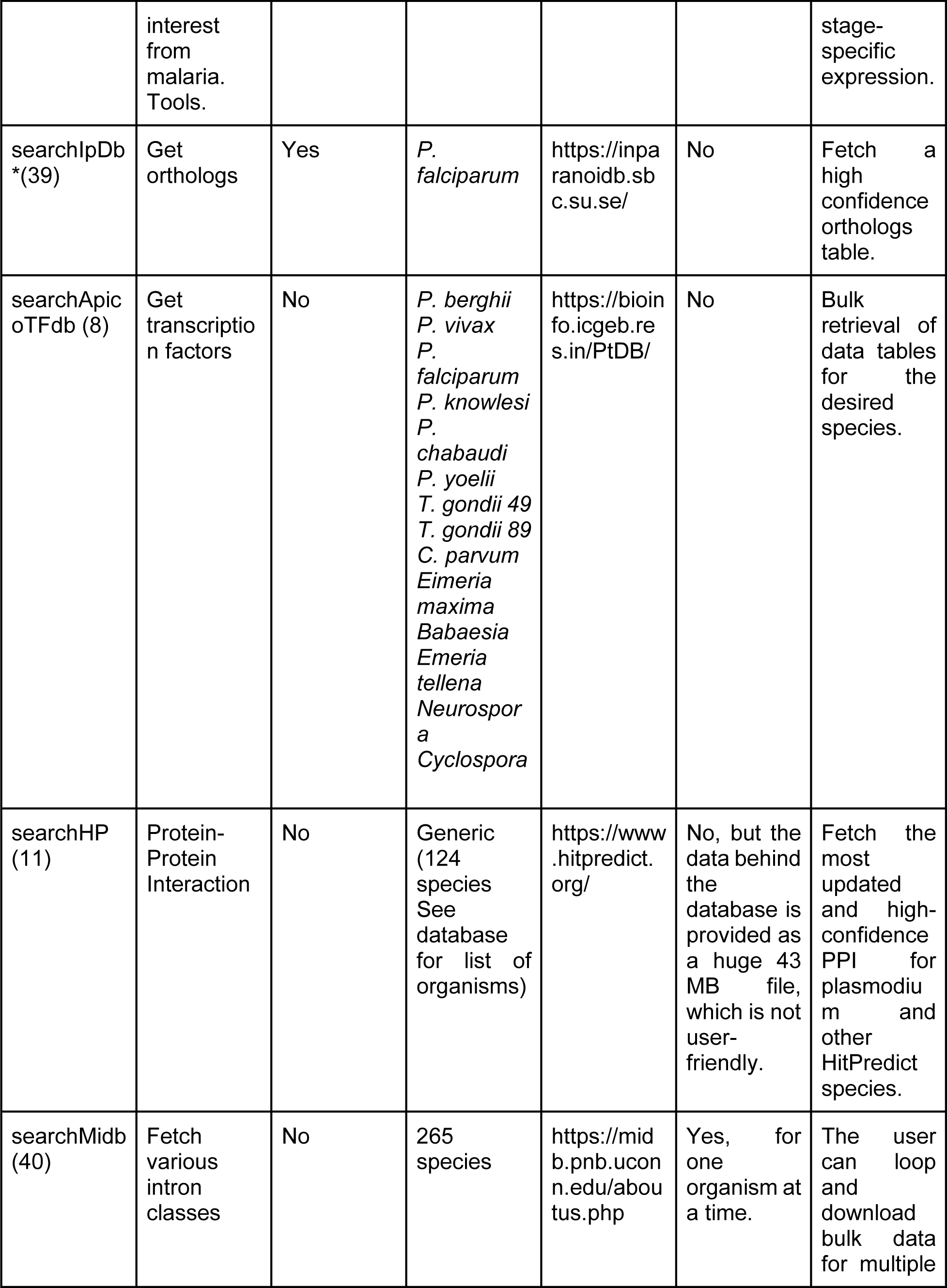
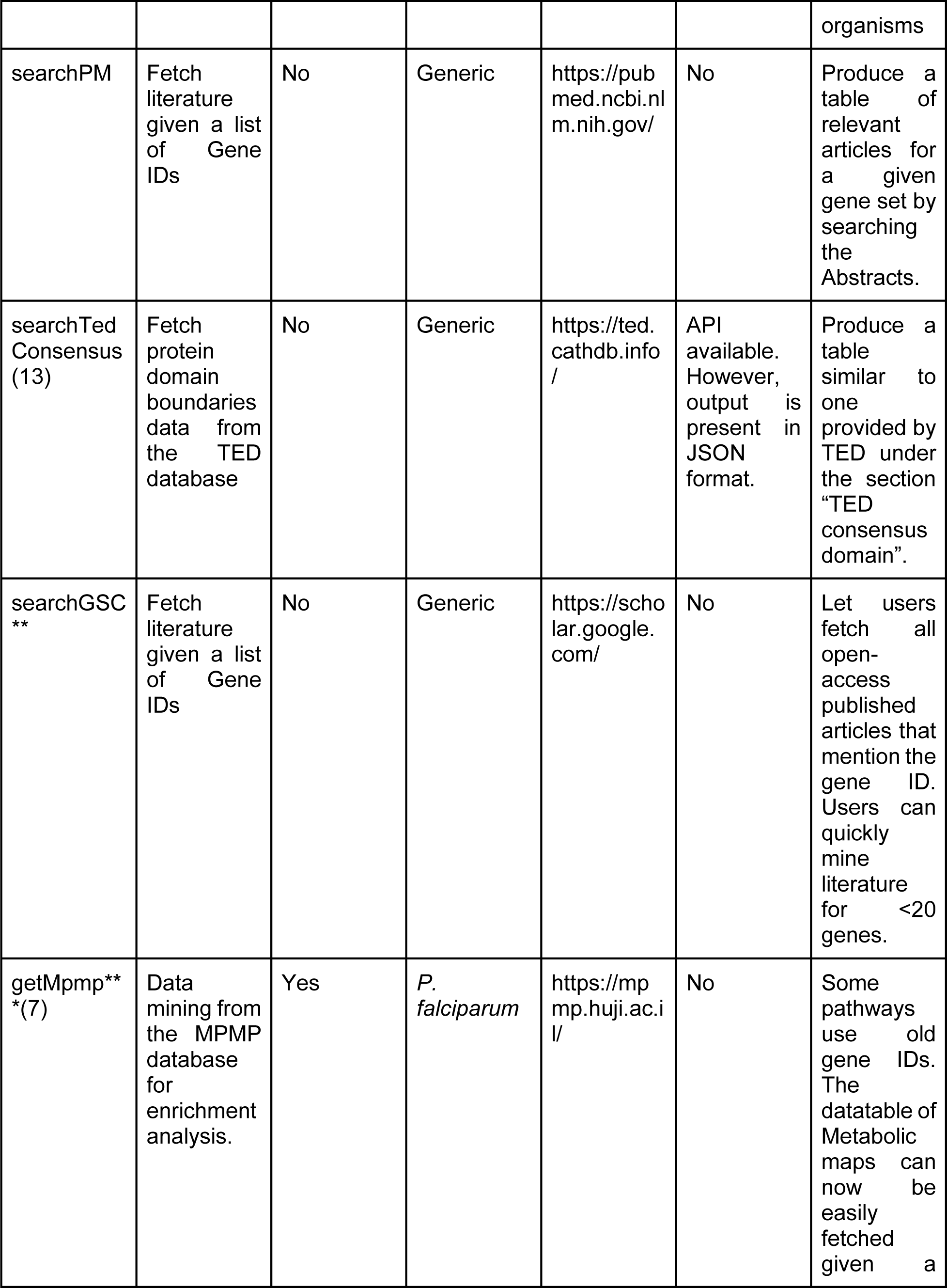
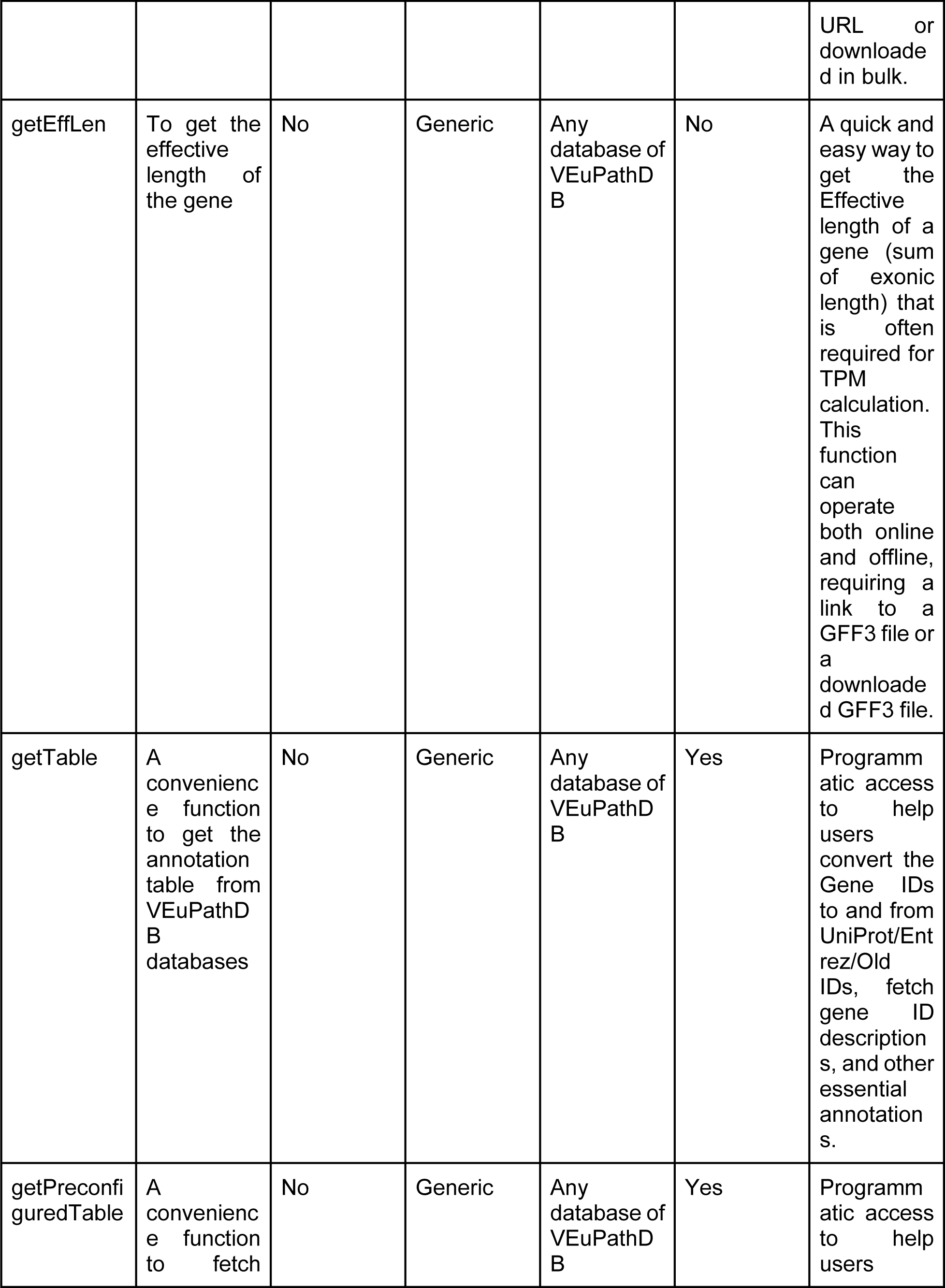
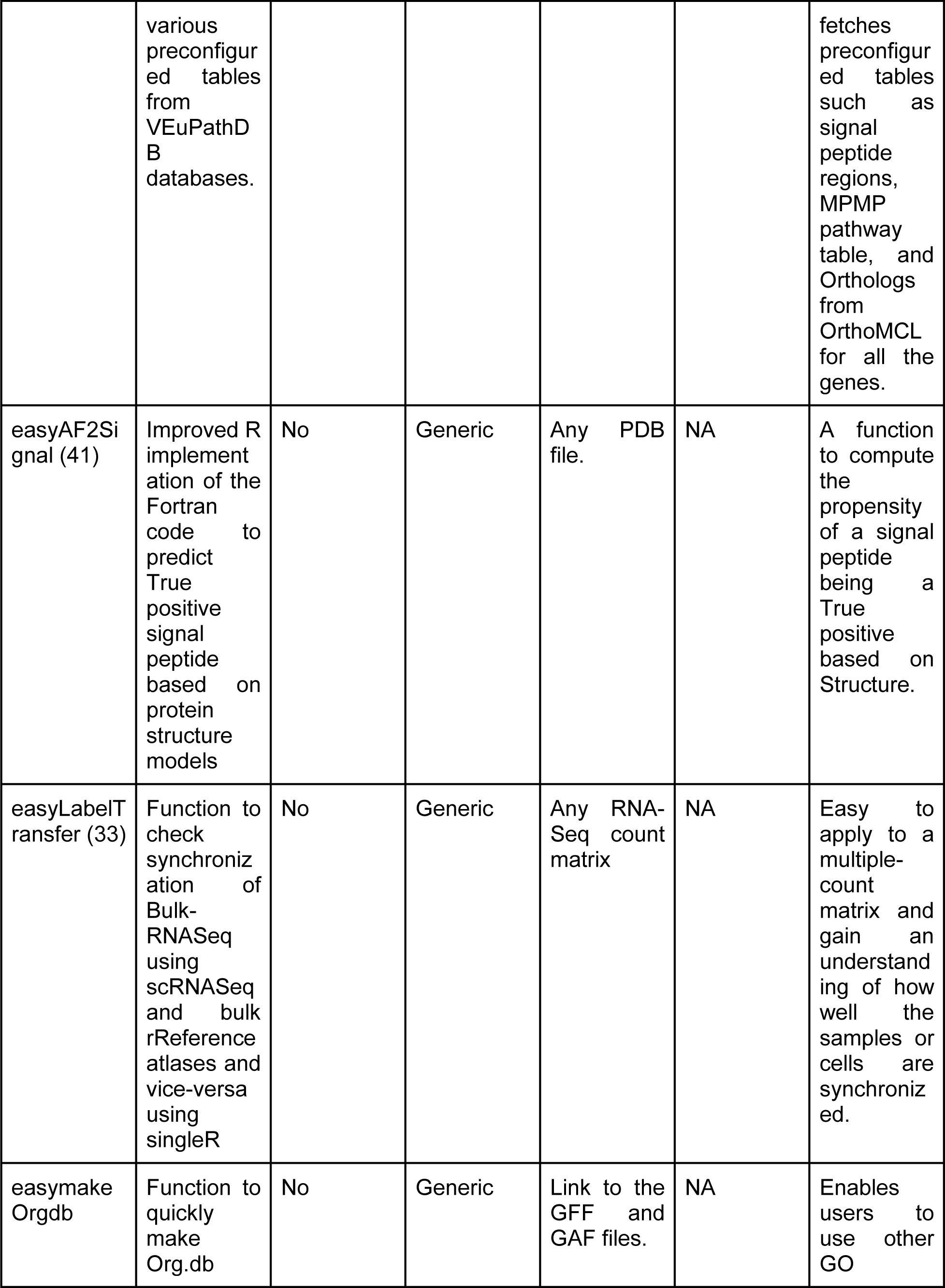
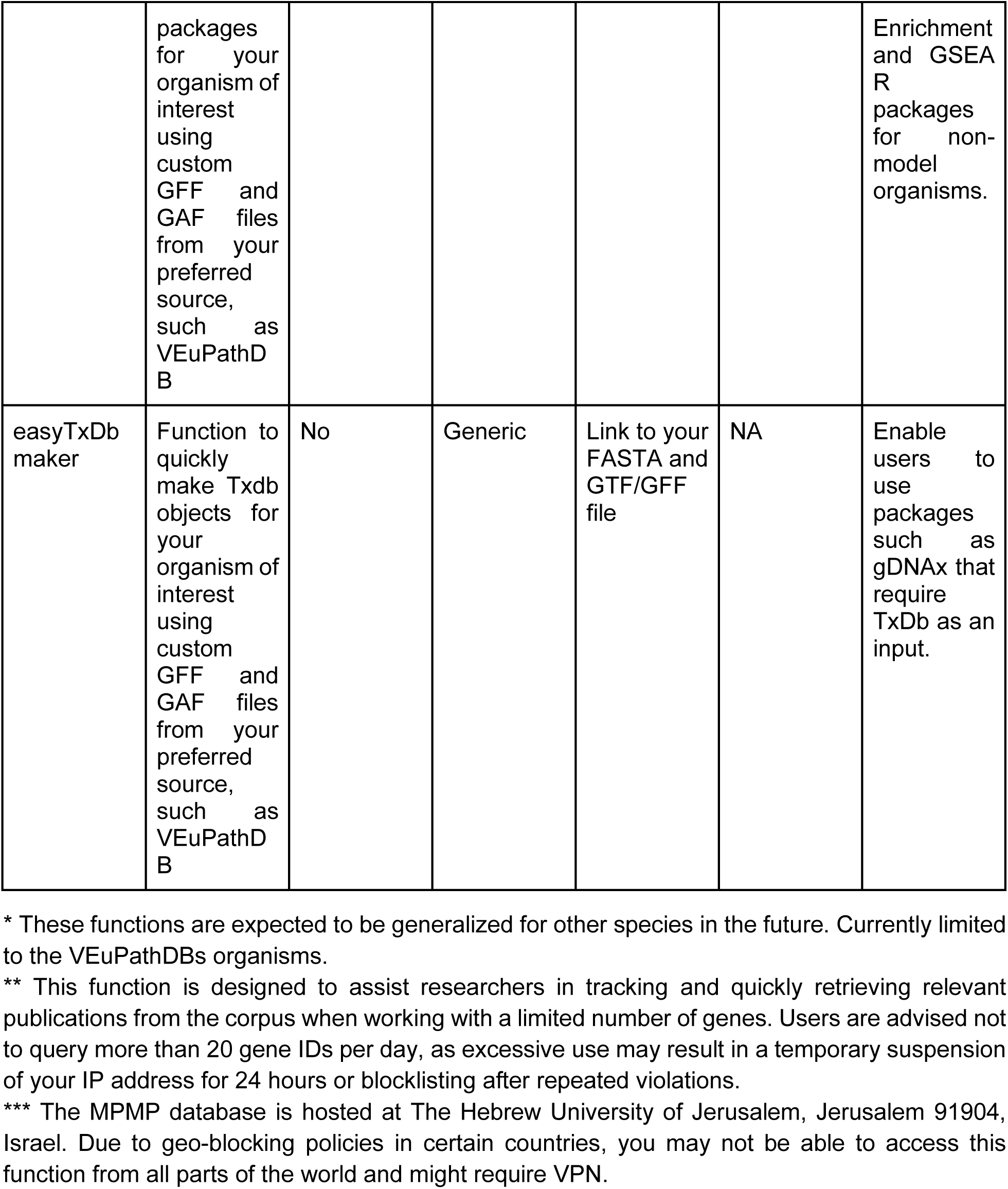
A list of select search and utility functions in the *plasmoRUtils* package. The search functions enable batch queries across multiple databases, while the utility functions simplify various downstream transcriptomics analyses.

*plasmoRUtils* utilizes several CRAN and Bioconductor packages in the background. These include data transformation, retrieval, and storage packages such as *rvest* (24), *dplyr* (25), *tidyr* (26), *easyPubMed*, *biomaRt* (22), *Biostrings* (27), *GenomeInfoDb*, *SingleCellExperiment* (28), *stringr*, *AnnotationForge*, *S4Vectors*, *tibble*, *table*, *GenomicFeatures* (29), *httr*, *readr*, *rlang*, *jsonlite*, *txdbmaker*, *IRanges*, *BiocGenerics*, *methods*, *janitor*, *magrittr*, *rtracklayer*, *plyr*, *purrr*, and *glue*. Visualization and documentation packages include *ggsci* (30), *drawProteins* (31), *scales*, *ggplot2*, *plotly*, *echarts4r*, *rmarkdown*, *knitr*, *BiocStyle*, *randomcoloR*, and *styler*. Finally, data analysis-specific packages include *topGO* (32), *SingleR* (33), *bio3d* (34), *scuttle*, *DESeq2* (35), *pathfindR* (21), *Seurat* (36), and *pRoloc* (37).

### AI Usage Declaration

The authors utilized OpenAI’s ChatGPT (GPT-4, accessed via ChatGPT Pro) and DeepSeek to assist in language editing. The authors reviewed and edited all AI-generated content to ensure accuracy and originality.

## 3: Use cases

### 3.1 Using the Search Function

Users can query various databases through different search functions (see Table 1). For example, *searchKipho* allows users to retrieve all kinases and phosphatases in *Plasmodium* species from the KiPho database (38), while *searchApicoTFdb* enables users to fetch transcription factor data for various *Apicomplexan* members (8). The *searchIpDb* function allows batch retrieval of orthologs from the InparanoidDB 9 database (39). Additionally, *searchPhPl* provides the ability to query multiple gene IDs to assess gene disruptability information. In some cases, users may also wish to obtain additional information, such as the reference and submitter details for phenotype and mutant phenotype reports. This information is stored in sub tables, which typically require manual access. However, *searchPhPl* enables users to quickly retrieve these tables in real-time, eliminating the need for manual effort and providing the latest, curated phenotype information from malaria experts (10). An accessory function called *easyPhplplottbl* generates Disruptability and Mutant Phenotype summary tables in R, mirroring the style of Phenoplasm visualizations (while using phenotype.txt files generated from the “Advanced Search” option of Phenoplasm).

Since keeping track of recent literature can be time-consuming, we also offer functions for searching relevant literature related to genes of interest using a list of gene IDs, that helps users stay informed on newly submitted research articles available through preprints. The two search functions available include: *searchGSC* (for accessing Google Scholar) and *searchPM* (for searching PubMed). Prior efforts have been made to assist researchers in keeping informed about. Several R packages developed by the ropensci community, such as *aRxiv* (https://github.com/ropensci/aRxiv, primarily for non-bio papers) and *medrxivr* (https://github.com/ropensci/medrxivr, for accessing articles from medarchive and bioarchive), aim to help with this. While *medrxivr* provides access to medRxiv and bioRxiv, users must create a snapshot of bioarchive data before using the function, which can be time-consuming, depending on the selected time range. For example, downloading data from October 2024 to January 2nd, 2025 (three months) took 16 minutes to retrieve 16,209 records. Since preprints on biorxiv, medrxiv, and other preprint platforms are indexed in Google Scholar (GS), we adopted a web-crawling strategy to fetch these articles, which proved to be faster using our custom *searchGSC* function. This function can also be used to gather an initial set of literature when working on a hypothetical protein by retrieving articles mentioning the gene ID of interest.

However, it is essential to note that while *searchGSC* can help track grey literature (articles not formally published by academic publishers but available as preprints), it will not retrieve articles that are not indexed in GS. Therefore, it should not be relied upon for comprehensive literature reviews (42). Additionally, there are limitations on the number of queries that can be made using this function (20 queries per gene per day, per IP address or host). Excessive use may result in the temporary blocking of the IP address.

### 3.2 Retrieving Preconfigured and User-Defined Annotations from VEuPathDBs

Bioinformaticians frequently face challenges related to gene ID conversion, particularly when working with legacy datasets (43). These tasks often involve mapping outdated gene IDs or converting between different ID formats, such as transcript, UniProt, or Entrez IDs, back to gene IDs, or replacing existing gene IDs with orthologous IDs from related apicomplexans. While VEuPathDB simplifies this process, ID conversion cannot always be automated or integrated into a pipeline when dealing with heterogeneous datasets, such as those with a mix of new gene IDs, gene symbols, and older IDs. This issue is particularly prevalent in resources such as Phenoplasm (with currently uses outdated mitochondrial gene IDs) and protein-protein interaction databases like MIIP, as well as databases like HitPredict, which provides PPI data using UniProt IDs, and BioCyc, which contains inconsistent mixes of old and current gene symbols and gene IDs.

While tools like the *AnnotationDbi::mapIds()* function in conjunction with the *Org.db* package or *biomaRt* could be used for these conversions, they have limitations. *Org.db* packages do not exist for all apicomplexan species, and *biomaRt* does not support all apicomplexans either (22). To address this gap, *plasmoRUtils* offers bioinformaticians the ability to quickly fetch the latest annotations and pre-configured tables, such as orthologs, MPMP pathways, and MetaCyc pathways, through "get" functions like *getPreconfiguredTable* and *getTable*. These functions enable users to rapidly retrieve extensive data tables from VEuPathDBs, which can then be utilized, along with custom coding, to map old gene IDs to new ones.

To streamline this process, especially since most gene IDs (such as PF3D7_XXXXX or TGME49_XXXXX for *P. falciparum* 3D7 genes and *Toxoplasma gondii* ME49 strain, respectively) are preferred over gene symbols (which are often absent), we developed a generic wrapper function, *toGeneid()*. This function builds on *getTable()* and enables users to quickly convert deprecated, Entrez, or UniProt IDs to new gene IDs, retrieve gene descriptions, and access other gene-related information such as GO terms, signal peptide sequences, and nearly any or all of the 90 fields or more available in the VEuPathDBs’ download tables. We evaluated the *getTable()* function by retrieving 90 fields (a large query) for all 5,791 genes from PlasmoDB—the query executed in 15.71 seconds, demonstrating efficient performance despite the high volume of data.

Additionally, the *getMPMP* function allows users to batch-fetch metabolic pathways and the genes involved in those pathways from the MPMP database for the first time. Users provide a set of URLs (obtained via the *data("listmpmp")* function), which are passed to *getMPMP()*. As the MPMP database does not consistently follow a clear data rendering structure (with some tables rendered as collapsible and others as static), the function attempts to identify the table type for each URL. For collapsible tables, which are more structured, the function returns all available columns (e.g., for http://mpmp.huji.ac.il/maps/gen_phosph.html, it returns columns such as EC Number and Formal annotation). For static tables (eg, http://mpmp.huji.ac.il/maps/HP1_enrich.html), where delimiters are inconsistently placed, only gene IDs and pathway names are returned.

### 3.3 Miscellaneous functions

In addition to data and annotation retrieval, *plasmoRUtils* also provides several wrapper functions to streamline standard transcriptomic downstream analyses using FASTA, GFF, and GAF files from VEuPathDBs. These functions, categorized as "easy" functions, include:

1. **Over-representation analysis (ORA):** VEuPathDBs offers web services for performing over-representation analysis; however, when the database is offline these become cumbersome making such analyses difficult and time-consuming (4,5). Moreover, VEuPathDB’s online services do not allow users to define custom background gene sets for Gene Ontology (GO) enrichment analysis. This is known to affect the over-representation results, such as when dealing with single-cell RNA-Seq where not all genes are successfully detected or if some genes were filtered before calling differentially expressed (DE) genes (44–47). We wrote a generic wrapper function called *easytopGO* that can help users perform ORA effortlessly (32). Given a set of differentially expressed genes (ranked or unranked), this wrapper function enables users to perform quick enrichment analysis, which can be plotted into publication-ready figures using the *easyGOPlot function*. Since the source of GO annotation also affects what users see in the ORA results, *easytopGO* allows flexibility to the users to either use Ensembl Protists GO annotations via BioMart or provide the GAF files from VEuPathDBs’. The package also provides functions to build Org.db packages using user-provided FASTA and GTF/GFF files, as well as the *easymakeOrgdb* function. The tar.gz file produced can then be easily installed and used with other GO enrichment R packages that utilize Orgdb objects, such as clusterProfiler (48). Another function, called easyTxDbmaker, can help users create TxDb objects using custom genomes and GTF/GFF files that are often required by packages such as *gDNAx* for assessing genomic DNA contamination.
2. **Compute TPM values:** Given a matrix of raw gene counts and effective gene lengths calculated by the *getEffLen* function, the *easyTPM* function can convert raw counts into TPM (Transcripts Per Million) values. The *getEffLen* function accounts for gene strand orientation, preventing the erroneous summing of overlapping exons in genes. The package also provides a plotting function called *easyExpPlot,* which enables users to create bubble plots or line plots for genes of interest or clusters of genes with similar expression patterns across different time points or sample types.
3. **Data Retrieval from MCA (Malaria Cell Atlas):** The *plasmoRUtils* package includes the *easyMCA* and *listMCA* functions, which provide direct access to malaria single-cell datasets from the Malaria Cell Atlas (MCA) (49). These functions enable users to fetch data sets without leaving R, making it easier to work with datasets on HPC machines or cloud platforms, such as Amazon Web Services (AWS). Users can view updated lists of available datasets using *listMCA*, and the output of *easyMCA* can be passed to the *easyLabelTransfer* function. This function enables the transfer of labels between bulk RNA-Seq datasets (such as time-series data) and single-cell RNA-Seq data, facilitating the integration of data from different sources. We also provide an easy-to-use wrapper function, *easyLabelTransfer*, based on the *singleR* package, to perform sample quality control. This function compares bulk time-series datasets with single-cell atlases, such as MCA, to verify sample stage/age (pf-ch10x-set4-biorxiv). As a user case, we applied this function to explore the time-point distribution of cells in the MCA, revealing an under-representation of time points (e.g., 20, 24, 28, and 32 hpi) (See Figure 2). In addition, we reanalyzed two *Toxoplasma* datasets obtained from (50,51) using the latest annotations from ToxoDB Release 68 for Toxoplasma and are providing them as reference atlases in the form of SingleCellExperiment objects (see Supplementary Materials for details).
4. **Signal Peptide fidelity assessment:** We offer a convenient function, *easyAF2Signal*, to diagnose potential false-positive signal peptides using SignalP predictions from the VEuPathDB database and AlphaFold2 structures, based on an observational study conducted by (41). Sanaboyana & Elcock et al. found that true N-terminal signal peptides (∼24-25 amino acids) are typically disengaged from the protein body and lack atomic contacts, and AlphaFold2 attempts to model them similarly. We developed an R equivalent with slight modifications to the Fortran code provided by the authors, which reports additional information, such as the number of residues remaining after pLDDT filtering. This is crucial because, if all the signal peptide residues have low pLDDT scores and are filtered out, no residues will remain to calculate contacts with the protein body, resulting in zero contacts. This could give a false impression that the first 25 amino acids are signal peptides, but it cannot be confirmed, as the zero-contact observed is due to no residues remaining after filtering, not because the signal peptide was disengaged from the protein body.
5. **NOISeq Annotation dataframe:** The easyNOISeqAnnot function is a convenience function that enables users to produce all the biological annotations required by the NOISeq package (52) to prepare the NOISeq object using user-provided GFF/GTF and FASTA files. QC functions provided by NOISeq function relies on this annotation to produce QC plots.
6. **PDB IDs to Uniprot ID Conversion:** When working with multi-chain protein complexes, researchers often need to map structural Chain IDs to Ensembl gene IDs. This conversion can be achieved through our package in two steps. The *pdb2uniprot* function can be used to process PDB entries (whether monomers or large complexes) into their constituent UniProt IDs by querying the PDBe API data, automatically handling all chains in the structure. These UniProt identifiers can then be seamlessly converted to gene IDs using the companion *toGeneid* function, enabling complete chain-to-gene mapping for further analyses.

**Figure 2:**
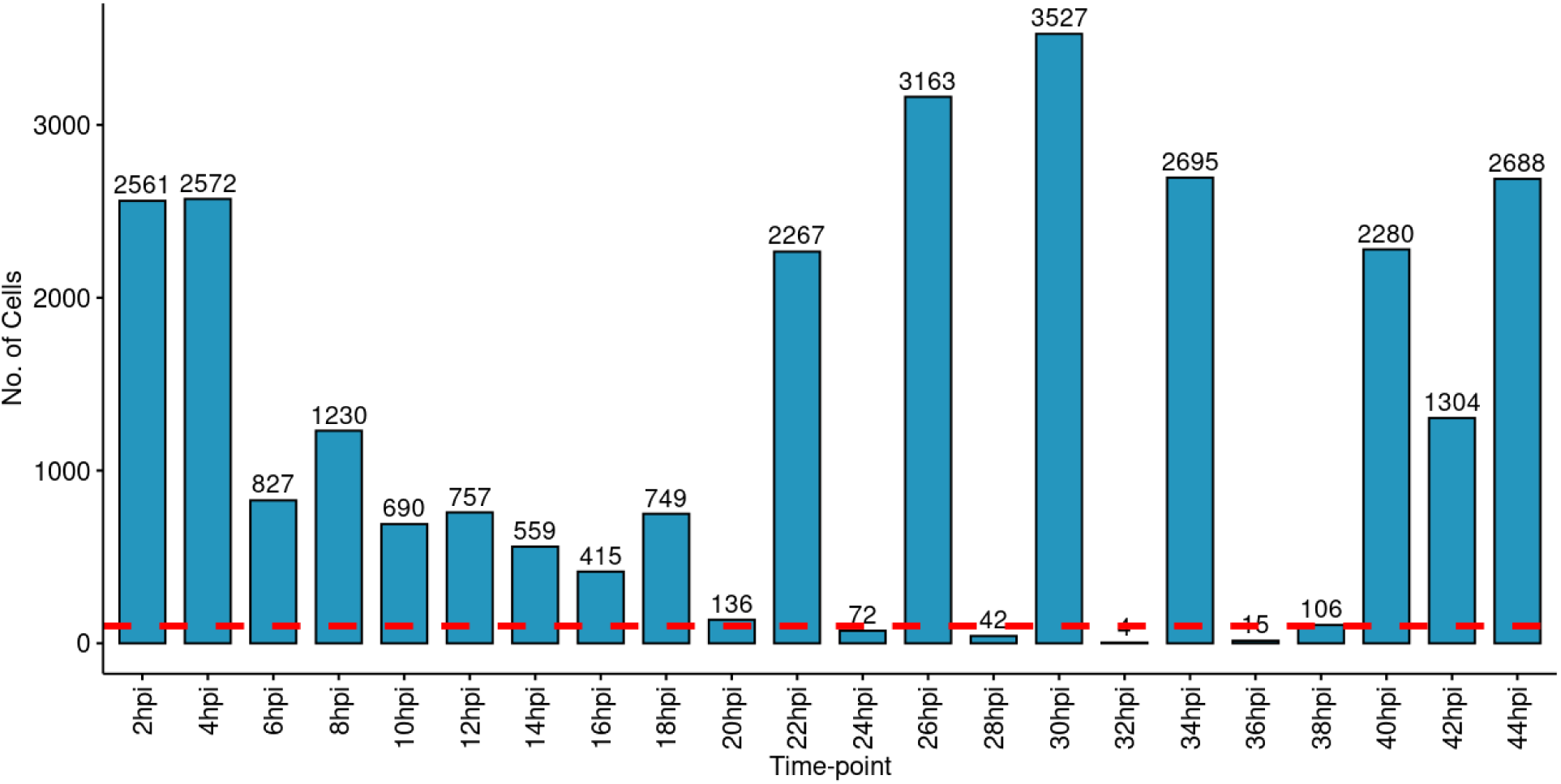
Analysis of cell distribution across various time points in the MCA, using the Bulk Time-Series dataset from Subudhi et al., 2020 and Toenhake et al., 2018. The red horizontal line indicates a cutoff of 100 cells. Notably, certain time points (e.g., 24, 28, 32, and 36 hpi) show a scarcity of cells in the malaria cell atlas i.e, <100 cells. Besides, in comparison to later time points (post 20 hpi), the early time point has a relatively smaller number of cells. These diagnostic plots can guide decisions to enrich the atlas with cells from previously underrepresented time-points/stages.

The aforementioned functions can also be integrated into pipelines, as illustrated in Figure 3. We demonstrate this using RNA sequencing data analysis as an example.

**Figure 3:**
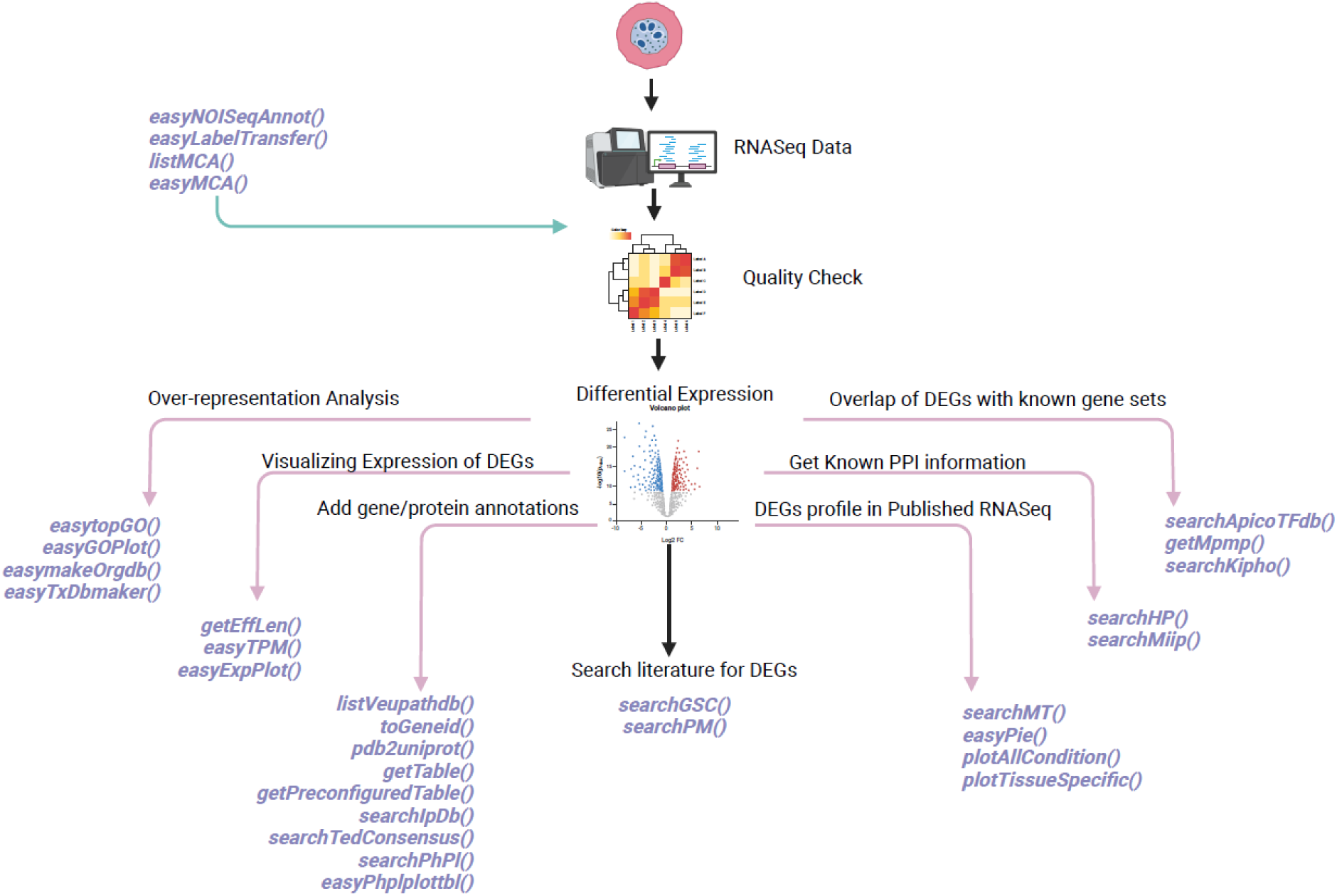
An exemplary overview of where the plasmoRUtils function can be used in the RNASeq data analysis workflow.

## 4. Discussion and Outlook

The *plasmoRUtils* is the first R package specifically designed to address several data access and transformation challenges central to Apicomplexan biology. By leveraging a combination of RESTful APIs and web-crawling techniques (when APIs are unavailable), the package enables streamlined retrieval and formatting of data from multiple disparate resources. In doing so, plasmoRUtils reduces the manual burden typically associated with working across heterogeneous databases and can serve as a generalizable template for interfacing with other complex or non-programmatically accessible datasets.

Our R package will continue to be updated biannually. We encourage users to visit our release page for updates on newly added or archived functions. As new databases featuring apicomplexan data emerge, such as *The Encyclopedia of Domains* (TED) (13), published in November 2024, we aim to provide access to them through *plasmoRUtils* (we recently added the *searchTedConsensus* function) (13). Future efforts would be directed to provide support for Pf7 dataset access and its exploration in R (53), inclusion of apicomplexan pathways from other sources such as LAMP (54) and protein interaction datasets, ortholog ID access and mapping using OrthoMCL-DB Release 7 (55), and additional wrapper and plotting functions.

## 5. Data Availability

As part of the *plasmoRUtils* package, we provide the most up-to-date data from the MPMP database, with the latest download performed on 28 August 2024. For legacy purposes, we also include MPMP pathway data (v2019), provided by PlasmoDB, for users who may require it. Additionally, we have transformed these datasets to facilitate their use with *pathfindR* for pathway enrichment analysis (21).

We have reanalyzed time-series data from (56) and (57) to assess time-point-based synchronization of cells and bulk samples. These datasets are provided as SummarizedExperiment objects within the package along with QC statistics (For details of reanalysis, read Supplementary Materials).

Furthermore, we have reanalyzed 10X scRNA-Seq data from (50,51), mapping all samples to the ME49 reference and using release 68 annotations, as significant changes in exon-intron boundaries has occurred since the publication of the original paper (See importance of reanalysis in vignette: https://rohit-satyam.github.io/plasmoRUtils/articles/Need_for_reanalysis.html and Supplementary Materials).

The data generated or reanalyzed during development of this package is included within the package and can be accessed via the *data()* function (58) for reproducibility.

## 6. Scope of improvement in *plasmoRUtils*

Although plasmoRUtils offers functions to access multiple databases, the successful retrieval of data depends on the maintenance and availability of those databases.

## Acknowledgments

We would like to express our gratitude to Kayenat Sheikh and other members of the Pathogen Genomics Laboratory at KAUST for sharing their experiences with the challenges of manual query and curation tasks, which highlighted areas that could benefit from automation. Finally, our sincere thanks go to Susanne Warrenfeltz, Omar S. Harb, Ulrike Boehme and Nupur Kittur from VEuPathDB team for providing extensive assistance in addressing our queries and providing us with data during the period when VEuPathDB went offline. We also thank the VEuPathDB developers and their team for providing access to their database via a REST API, around which we built some of our functions, and for keeping it freely accessible. Lastly, we express our gratitude to Yuan Xue (original author of the SMARTseq2 scRNASeq Toxo atlas) for providing valuable input during data reanalysis.

## Note

This work aims to provide CLI users and bioinformaticians with seamless and efficient access to VEuPathDB, its constituent databases, and other resources within their pipelines. However, this initiative is not intended to discourage users from subscribing to the database. VEuPathDB remains a critical resource for parasite research, and the continued utility of some of the functions in this package relies on community support and database maintenance. We strongly encourage all users who benefit from VEuPathDB to support the database team and, in turn, the database itself by subscribing. We also encourage users to cite VEuPathDB or relevant databases when citing the use of our R package.

## Availability of source code and requirements

The package plasmoRUtils has been tested on the following OS:

1. Windows: AMD Ryzen 7 6800H with Radeon Graphics with installed RAM 16.0 GB (15.2 GB usable) 64-bit operating system, x64-based processor
2. MacOS: Macbook Pro, M1 2020, Sonoma 14.2.1 with RAM 16.0 GB
3. Ubuntu 20.04.6 LTS: Intel® Xeon(R) Gold 6330 CPU 2.00GHz × 112

Programming language: R > 4.0

## Contributions

RS: Conception, development, data analysis, and manuscript preparation; AM and DGC: Package testing, documentation, test runs, and package maintenance and deployment; AKS: Ideation of various utility functions and providing test data and use cases for them, and function improvement. RPS: Helped with SmartSeq2 data analysis; AP: Funding, Manuscript preparation, revision, structuring, and submission.

## Ethical approval

Not applicable.

## Consent for publication

Not applicable.

## Supplementary Materials

### Reanalyzed Datasets

We reanalyzed the Single Cell RNA-Seq dataset of *Toxoplasma gondii* from two major studies and intend to make this data available for community use and as a ready-to-use reference atlas for the *easyLabelTransfer()* function. This effort aims to make the reanalyzed single-cell datasets available, using the latest annotations from VEuPathDB, as the intron-exon boundaries in the database are continually evolving. We observed that the reanalysis with improved annotation leads to an increase in the number of genes detected per cell, mean reads per cell, and could rescue a greater number of cells. The following methodology was used for reanalysis.

#### 1. Preprocessing

For preprocessing (51), we obtained the SMART-Seq2 data for ME49 and RH strains using fasterq-dump 3.1.1. The reference genome and annotation were obtained from the ToxoDB database (release 68) and indexed using *--sjdbOverhang 99* for all samples, except for the RH 96 well plate, where *--sjdbOverhang 70* was used. ERCC genes were also added to the annotation and were obtained from https://github.com/DeplanckeLab/SmartSeq2/blob/master/ERCC92.fa. First ribodepletion was carried out using Ribodetector 0.3.1 using a more stringent mode where we use parameter *-e norrna* to get rid of moderate to high confidence rRNA reads, as the authors reported high rRNA contamination. The RNAgrinder nextflow workflow (https://github.com/Rohit-Satyam/RNAgrinder) was then modified to analyse the SMARTSeq2 data. The reads were then aligned to the genome using STAR solo v2.7.11a with the following modifications in the RNAgrinder code. *-- outFilterMatchNminOverLread 0.3 --outFilterScoreMinOverLread 0.3 --soloType SmartSeq -- outSAMstrandField intronMotif --soloUMIdedup Exact NoDedup --soloStrand Unstranded -- soloFeatures Gene GeneFull SJ*. The genes were quantified using featureCounts v2.0.8 with *-- fraction and -M -s 0 --countReadPairs* flags to account for multimapping reads, similar to the author’s approach. This was followed by the use of the *round()* function to obtain absolute counts. To identify poor-quality cells, we first load the counts as a SingleCellExperiment object and leverage ERCC spike-in genes with *scater::quickPerCellQC()* to identify poor-quality cells. This led us to recover 641 cells from ME49day0, 998 from ME49day3, 615 from RH384well, and 253 cells from RH96well. The cells were converted to a Seurat object. For (50), we downloaded the 10X 3’ v3 data from SRA using fastq-dump and fed it to CellRanger v7.1.0 with intron mode. We recovered 12,741 cells (4,700 more cells than reported by the authors) with a mean of 6,217 reads per cell and a median of 537 expressed genes per cell. We load the filtered counts as a Seurat object in R using the same filtering thresholds as used by the authors (*min.cells = 5, min.features = 100*). The QC information has been provided in the metadata.

#### 2. Reannotation of reanalyzed data

We created an integrated reference of ME49 and RH Strain by borrowing the metadata and raw counts provided by these studies using the CCAIntegration method of the *IntegrateLayers()* function and evaluated whether the integration was proper by examining the UMAP (See below). We then used the *TransferData()* function to transfer the labels from this joint atlas back to the reanalyzed datasets (See Supplementary Figures 1, 2, and 3).

The data has been provided as SingleCellExperiment objects with the package.

**Supplementary Figure 1:**
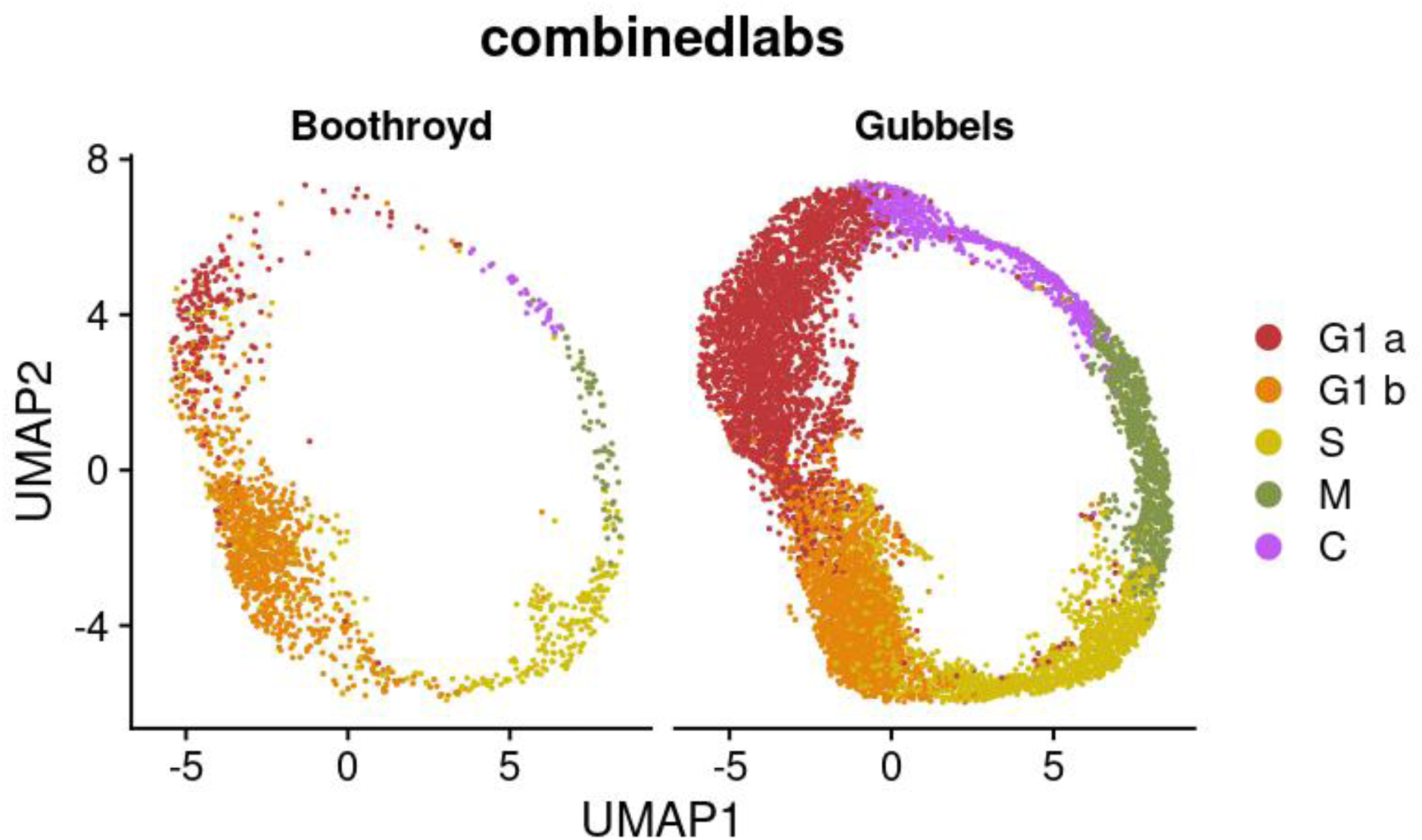
Integration of SmartSeq2 (ME49) and 10X (RH strain) scRNASeq dataset of *Toxoplasma gondii (old references)*.

**Supplementary Figure 2:**
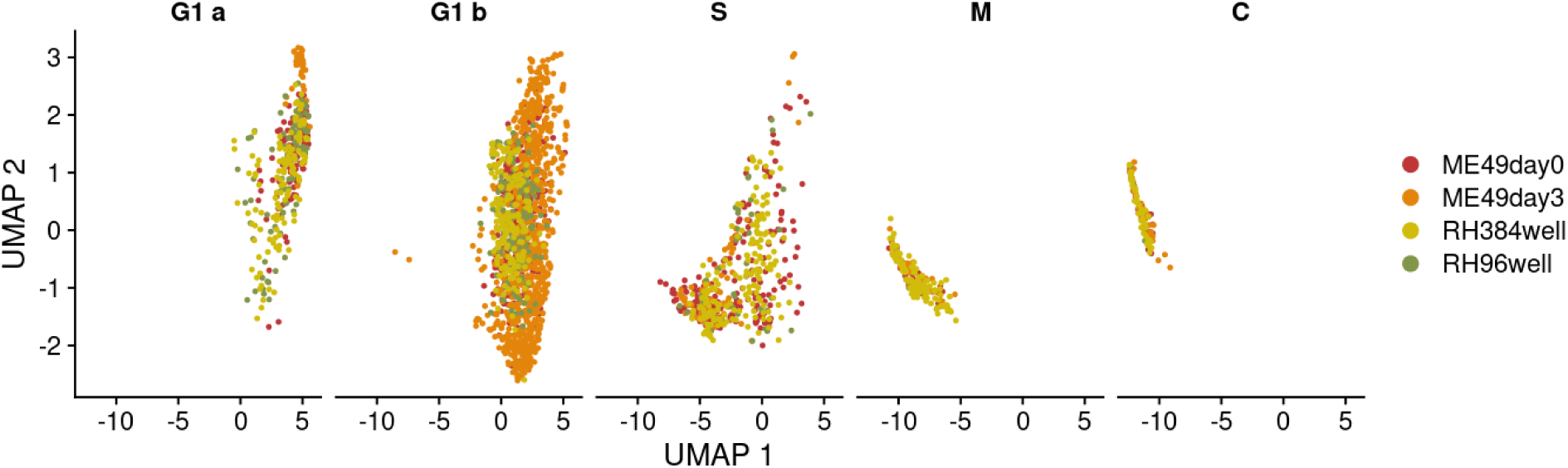
Successful label transfer from old reference atlas to reanalyzed SmartSeq2 (shown as facets to show proper label assignment) samples from Xue et al, 2020, and 10x. UMAP shows proper label transfer. For 10x since barcode information can be used to check the accuracy of label transfer, we saw 95% of barcodes have been assigned the same label.

#### 3. Bulk RNA-Seq analysis of Time-Series dataset

We reanalysed two published plasmodium bulk RNASeq datasets from available from (Subudhi et al., 2020) and (Toenhake et al., 2018) using *Plasmodium falciparum* 3D7 annotation from the current release (v68). We used the RNAgrinder nextflow pipeline with *--sjdbOverhang 99 and --sjdbOverhang 74,* respectively to index the genomes and perform alignment using the STAR aligner. The featureCounts was used with similar parameters (-M -C --fraction) to account for multimapping reads.

## References

1. Bhatia A, Pruthi P, Chakraborty I, Shukla N, Narayan J. getENRICH: a tool for the gene and pathway enrichment analysis of non-model organisms. Bioinforma Adv. 2025;5(1).

2. Amos B, Aurrecoechea C, Barba M, Barreto A, Basenko EY, Bażant W, et al. VEuPathDB: The eukaryotic pathogen, vector and host bioinformatics resource center. Nucleic Acids Res. 2022;

3. Aurrecoechea C, Barreto A, Basenko EY, Brestelli J, Brunk BP, Cade S, et al. EuPathDB: The eukaryotic pathogen genomics database resource. Nucleic Acids Res. 2017;

4. Fernandez-Prada C, Moretti NS, do Monte-Neto RL. Critical loss: the effects of VEuPathDB defunding on global health. The Lancet Microbe. 2024;1–3.

5. Keroack C, Cosentino RO, Teixeira TL, Lansink L, Larcombe SD, Davidge B, et al. Scientists swarm into Woods Hole for the 10th Kinetoplastid Molecular Cell Biology Meeting. Trends Parasitol [Internet]. 2024;40(12):1057–62. Available from: 10.1016/j.pt.2024.10.009

6. Tan QW, Mutwil M. Malaria.tools-comparative genomic and transcriptomic database for Plasmodium species. Nucleic Acids Res. 2020;

7. Ginsburg H, Abdel-Haleem AM. Malaria Parasite Metabolic Pathways (MPMP) Upgraded with Targeted Chemical Compounds. Trends in Parasitology. 2016.

8. Sardar R, Kaushik A, Pandey R, Mohmmed A, Ali S, Gupta D. ApicoTFdb: The comprehensive web repository of apicomplexan transcription factors and transcription-associated co-factors. Database. 2019;

9. Bhattacharyya M, Chakrabarti S. Identification of important interacting proteins (IIPs) in Plasmodium falciparum using large-scale interaction network analysis and in-silico knock-out studies. Malar J. 2015;

10. Sanderson T, Rayner JC. PhenoPlasm: A database of disruption phenotypes for malaria parasite genes. Wellcome Open Research. 2017.

11. López Y, Nakai K, Patil A. HitPredict version 4: Comprehensive reliability scoring of physical protein-protein interactions from more than 100 species. Database. 2015;

12. Patil A, Nakai K, Nakamura H. HitPredict: A database of quality assessed protein-protein interactions in nine species. Nucleic Acids Res. 2011;

13. Lau AM, Bordin N, Kandathil SM, Sillitoe I, Waman VP, Wells J, et al. Exploring structural diversity across the protein universe with The Encyclopedia of Domains. Science (80- ) [Internet]. 2024;4946:2024.03.18.585509. Available from: https://www.biorxiv.org/content/10.1101/2024.03.18.585509v2 https://www.biorxiv.org/content/10.1101/2024.03.18.585509v2.abstract

14. Lasonder E, Green JL, Grainger M, Langsley G, Holder AA. Extensive differential protein phosphorylation as intraerythrocytic Plasmodium falciparum schizonts develop into extracellular invasive merozoites. Proteomics. 2015;

15. Mok S, Stokes BH, Gnädig NF, Ross LS, Yeo T, Amaratunga C, et al. Artemisinin-resistant K13 mutations rewire Plasmodium falciparum’s intra-erythrocytic metabolic program to enhance survival. Nat Commun. 2021;

16. Tewari SG, Kwan B, Elahi R, Rajaram K, Reifman J, Prigge ST, et al. Metabolic adjustments of blood-stage Plasmodium falciparum in response to sublethal pyrazoleamide exposure. Sci Rep. 2022;

17. Wang C, Dong Y, Li C, Oberstaller J, Zhang M, Gibbons J, et al. MalariaSED: a deep learning framework to decipher the regulatory contributions of noncoding variants in malaria parasites. Genome Biol. 2023;

18. Shang X, Wang C, Shen L, Sheng F, He X, Wang F, et al. PfAP2-EXP2, an Essential Transcription Factor for the Intraerythrocytic Development of Plasmodium falciparum. Front Cell Dev Biol. 2022;

19. Aurrecoechea C, Brestelli J, Brunk BP, Dommer J, Fischer S, Gajria B, et al. PlasmoDB: A functional genomic database for malaria parasites. Nucleic Acids Res. 2009;

20. Stoeckert CJ, Fischer S, Kissinger JC, Heiges M, Aurrecoechea C, Gajria B, et al. PlasmoDB v5: new looks, new genomes. Trends in Parasitology. 2006.

21. Ulgen E, Ozisik O, Sezerman OU. PathfindR: An R package for comprehensive identification of enriched pathways in omics data through active subnetworks. Front Genet. 2019;

22. Durinck S, Spellman PT, Birney E, Huber W. Mapping identifiers for the integration of genomic datasets with the R/ Bioconductor package biomaRt. Nat Protoc. 2009;

23. Lin S, Scott D. Webscraping with rvest. In: Hands-On Data Science for Librarians. 2023.

24. Wickham H. rvest: Easily Harvest (Scrape) Web Pages R package version 1.0.4. 2024; Available from: https://rvest.tidyverse.org/

25. Wickham H, Francois R. The dplyr package. R Core Team. 2016;

26. Hadley Wickham and Davis Vaughan and Maximilian Girlich. Tidy Messy Data • tidyr. R package version. 2024.

27. Pagés H, Aboyoun P, Gentleman R, DebRoy S. Biostrings: Efficient manipulation of biological strings. R package version 2.58.0. 2020.

28. Lun A, Risso D, Korthauer K. {SingleCellExperiment}: {S4} classes for single cell data. R Packag version. 2018;

29. Carlson AM, Pagès H, Aboyoun P, Falcon S, Morgan M, Sarkar D, et al. Package ‘GenomicFeatures.’ 2021;

30. Xiao N. ggsci: Scientific Journal and Sci-Fi Themed Color Palettes for “ggplot2”. R package version 2.9. https://CRAN.R-project.org/package=ggsci. R package version 2.7. 2018.

31. Brennan P. DrawProteins: A bioconductor/R package for reproducible and programmatic generation of protein schematics. F1000Research. 2018;

32. Rahnenfuhrer AA. Bioconductor. 2022. Bioconductor - topGO.

33. Aran D, Looney AP, Liu L, Wu E, Fong V, Hsu A, et al. Reference-based analysis of lung single-cell sequencing reveals a transitional profibrotic macrophage. Nat Immunol. 2019;

34. Grant BJ, Skjærven L, Yao XQ. The Bio3D packages for structural bioinformatics. Protein Sci. 2021;

35. Love MI, Huber W, Anders S. Moderated estimation of fold change and dispersion for RNA-seq data with DESeq2. Genome Biol. 2014;

36. Hao Y, Stuart T, Kowalski MH, Choudhary S, Hoffman P, Hartman A, et al. Dictionary learning for integrative, multimodal and scalable single-cell analysis. Nat Biotechnol. 2024;

37. Crook OM, Breckels LM, Lilley KS, Kirk PDW, Gatto L. A Bioconductor workflow for the Bayesian analysis of spatial proteomics. F1000Research. 2019;

38. Pandey R, Kumar P, Gupta D. KiPho: malaria parasite kinome and phosphatome portal. Database (Oxford). 2017;

39. Persson E, Sonnhammer ELL. InParanoiDB 9: Ortholog Groups for Protein Domains and Full-Length Proteins. J Mol Biol. 2023;

40. Moulton JK, Wiegmann BM. Evolution and phylogenetic utility of CAD (rudimentary) among Mesozoic-aged Eremoneuran Diptera (Insecta). Mol Phylogenet Evol. 2004;

41. Sanaboyana VR, Elcock AH. Improving Signal and Transit Peptide Predictions Using AlphaFold2-predicted Protein Structures. J Mol Biol. 2024;

42. Haddaway NR, Collins AM, Coughlin D, Kirk S. The role of google scholar in evidence reviews and its applicability to grey literature searching. PLoS One. 2015;

43. Ikeda S, Ono H, Ohta T, Chiba H, Naito Y, Moriya Y, et al. TogoID: An exploratory ID converter to bridge biological datasets. Bioinformatics. 2022;

44. Geistlinger L, Csaba G, Santarelli M, Ramos M, Schiffer L, Turaga N, et al. Toward a gold standard for benchmarking gene set enrichment analysis. Brief Bioinform. 2021;

45. Reimand J, Isserlin R, Voisin V, Kucera M, Tannus-Lopes C, Rostamianfar A, et al. Pathway enrichment analysis and visualization of omics data using g:Profiler, GSEA, Cytoscape and EnrichmentMap. Nat Protoc. 2019;

46. Timmons JA, Szkop KJ, Gallagher IJ. Multiple sources of bias confound functional enrichment analysis of global -omics data. Genome Biol. 2015;

47. Ziemann M, Schroeter B, Bora A. Two subtle problems with overrepresentation analysis. Bioinforma Adv. 2024;4(1).

48. Yu G, Wang LG, Han Y, He QY. ClusterProfiler: An R package for comparing biological themes among gene clusters. Omi A J Integr Biol. 2012;

49. Howick VM, Russell AJC, Andrews T, Heaton H, Reid AJ, Natarajan K, et al. The malaria cell atlas: Single parasite transcriptomes across the complete Plasmodium life cycle. Science (80- ). 2019;

50. Lou J, Rezvani Y, Arriojas A, Wu Y, Shankar N, Degras D, et al. Single cell expression and chromatin accessibility of the Toxoplasma gondii lytic cycle identifies AP2XII-8 as an essential ribosome regulon driver. Nat Commun [Internet]. 2024;15(1). Available from: 10.1038/s41467-024-51011-7

51. Xue Y, Theisen TC, Rastogi S, Ferrel A, Quake SR, Boothroyd JC. A single-parasite transcriptional atlas of toxoplasma gondii reveals novel control of antigen expression. Elife. 2020;

52. Tarazona S, Furió-Tarí P, Turrà D, Di Pietro A, Nueda MJ, Ferrer A, et al. Data quality aware analysis of differential expression in RNA-seq with NOISeq R/Bioc package. Nucleic Acids Res. 2015;

53. Abdel Hamid MM, Abdelraheem MH, Acheampong DO, Ahouidi A, Ali M, Almagro-Garcia J, et al. Pf7: an open dataset of Plasmodium falciparum genome variation in 20,000 worldwide samples. Wellcome Open Res. 2023;

54. Shanmugasundram A, Gonzalez-Galarza FF, Wastling JM, Vasieva O, Jones AR. Library of Apicomplexan Metabolic Pathways: A manually curated database for metabolic pathways of apicomplexan parasites. Nucleic Acids Res. 2013;

55. Fischer S, Brunk BP, Chen F, Gao X, Harb OS, Iodice JB, et al. Using OrthoMCL to assign proteins to OrthoMCL-DB groups or to cluster proteomes into new ortholog groups. Curr Protoc Bioinforma. 2011;

56. Subudhi AK, O’Donnell AJ, Ramaprasad A, Abkallo HM, Kaushik A, Ansari HR, et al. Malaria parasites regulate intra-erythrocytic development duration via serpentine receptor 10 to coordinate with host rhythms. Nat Commun. 2020;

57. Toenhake CG, Fraschka SAK, Vijayabaskar MS, Westhead DR, van Heeringen SJ, Bártfai R. Chromatin Accessibility-Based Characterization of the Gene Regulatory Network Underlying Plasmodium falciparum Blood-Stage Development. Cell Host Microbe. 2018;

58. Attwood TK, Agit B, Ellis LBM. Longevity of Biological Databases. EMBnet.journal. 2015;

